# Human milk miRNAs respond to infections of the mother and infant during breastfeeding

**DOI:** 10.1101/2025.01.07.631768

**Authors:** Mohammed Alsaweed, Mezyndra Badsha, Ching Tat Lai, Donna T. Geddes, Foteini Kakulas

## Abstract

miRNAs have been recently discovered in different mammals’ milk. Specifically, human milk (HM) is highly rich in miRNAs, with differential expression amongst its fractions including cells, fat, and skim milk. Various factors, such as the stage of lactation or milk removal during breastfeeding have been shown to influence the miRNA content of HM. Here, we sought to determine the effect of maternal and/or infant infection on the miRNA profile of HM cell and fat fractions analyzed using next generation sequencing. Breastfeeding mother/infant dyads (n=18) were followed during one or more infection episodes as well as upon recovery. HM cells and fat together contained 1,780 known miRNA species, which is the highest number of known miRNAs assayed in human body fluids to date. In addition, 592 novel miRNAs were predicted, of which 95 were of high confidence. Comparisons between samples collected when the participants were healthy and when infected yielded 453 differentially expressed (p<0.05) known miRNAs. Of these, 70 were highly expressed and differentially regulated during infection, with 62 upregulated and 8 downregulated known miRNAs during infection. Most of the highly and differentially expressed miRNAs are known to play critical roles in immunity and immune system development. These findings support the use of HM miRNAs as biomarkers of the health status of the lactating breast and the breastfeeding mother/infant dyad.

## INTRODUCTION

A plethora of immune factors are transferred from the mother to the infant via human milk (HM) [1,2], providing protection against infections as well as signals that program the development of the infant’s own defenses [3–5]. HM-associated immune-protection is stronger during exclusive breastfeeding [6], which is recommended for the first six months after birth [7,8], with breastfed infants being documented to experience lower risks of infection and development of infectious diseases compared to their formula-fed counterparts [2,9]. These protective effects of HM are mediated by both cellular (leukocytes) and molecular (immunoglobulins, lactoferrin, lysozyme, etc.) components, which have been shown to rapidly respond to infections of the mother and/or the infant, and to be involved in the protection of both the mammary gland and infant from infection [6,10–13]. Very recently, miRNAs were identified as a new component abundant in HM, with immunomodulatory properties [14,15].

miRNAs are small (typically 22 nucleotides in length), non-coding RNA molecules regulating gene expression at the post-transcriptional level [16]. They function in the control of protein synthesis in numerous biological processes, including cell proliferation [17], apoptosis [18], cell differentiation [19], tissue development [20], immune response [21], and many others [22]. Specifically in relation to their involvement in immunity, miRNAs participate in the regulation of leukocyte function, such as the B lymphocytes [23,24], and the control of the adaptive and innate immunity [25,26]. miRNAs have been profiled using deep sequencing in HM cells [27–29], fat [27,29,30], skim milk [15], and exosomes[30], with the cell fraction of HM being the richest in miRNA amongst its other fractions, followed by HM fat [22,27–29]. These recent studies have suggested that miRNAs are very abundant in HM compared to other human body fluids [14,22,28]. This may partially be due to the endogenous synthesis of the majority of milk miRNAs in the mammary gland, which has been supported by both human and animal studies [27–29,31]. Further, HM miRNAs have been suggested to play unique roles in the mammary gland and for the infant, participating in the synthesis of milk components [29], and responding to milk removal during breastfeeding [29] as well as changing with the stage of lactation in response to the changing infant needs [28]. In addition, HM miRNAs have been suggested to protect the infant from pathogens and program the developing infant’s immune system and defense mechanisms [15,22,32].

Studies on the immunomodulatory properties of milk miRNAs are scarce and have primarily used animal models. Recently, bovine milk miRNAs have been examined during infection of the udder [33–35]. Differential miRNA expression and upregulation or downregulation of specific miRNA species was reported in response to *Staphylococcus aureus* and *Streptococcus uberis* infection [33,34], suggesting that miRNAs are key amplifiers of monocytic inflammatory response networks [34]. A commonly examined human milk immune-related miRNA is miR-150, which increases in B and T lymphocytes in response to various bacterial and viral infections [36,37]. Further, miR-181a, miR- 155 and miR-146a, found also in human milk, have been found to play critical roles in T and B cell differentiation and activation [38,39]. In innate immunity, some miRNAs such as miR-146a, which are rich in human milk, have been identified as upregulators of human monocytes stimulated with strong bacterial infection [40]. Interestingly, miRNAs encapsulated in exosomes or other milk-contained microvesicles have shown high stability to different harsh environments and conditions to capability features and play roles in the extracellular microenvironment, especially in body fluids, being involved in critical functions of immune response through intercellular communication [15,32,41]. Hence, it has been proposed that milk miRNAs, including those of HM, may serve as important biomarkers of infections of the mammary gland as well as to monitor the response to treatment and recovery [22,33,42,43]. In this study, we aimed to shed light into the HM miRNA response to infections of the breastfeeding dyad by profiling the miRNA composition of HM using Solexa Sequencing and performing qRT-PCR for specific immune-related miRNAs in different milk fractions (cells, fat, and skim milk) during both healthy conditions and natural occurring infections of the breastfeeding mother and infant.

## MATERIALS AND METHODS

### Ethics statement and sample collection

This study was approved by the Human Research Ethics Committee of The University of Western Australia (Ethics Number: RA/4/1/4397). All participating mothers provided written informed consent. The demographic characteristics of the participating mother/infant dyads are shown in Table 1. Breastfeeding mothers (n=18) were recruited during healthy and natural occurring infection periods. Freshly expressed HM samples (22-158 mL; total samples collected n=40) were collected from each mother whilst she and her infant were both healthy, and participants were followed for collection of further HM samples if either the mother or the infant or both experienced an infection **(Table 1)**. On some occasions, more than one infections were sampled from a specific dyad. Infections types included: common cold/flu (n=10 cases), mastitis (n=3 cases), Recurring Ear infection (n=3 cases), Recurring infection including Mother Ulcerative colitis/sweets syndrome, Hashimoto/baby well (n=2 cases), Gastro infection (n=3 cases), Toe Infection (n=1 case), and cracked nipples (n=1 case). Samples were collected aseptically using an electric breast pump (Medela AG, Baar, Switzerland), and were immediately transported to the laboratory for analyses.

**Table 1.** The range (mean ± standard deviation, S.D.) of the mothers demographic characteristics, HM cell content, HM fat content, HM RNA content and quality in all HM samples including all HM fractions (n=117), samples collected during infection (n=63), and samples collected during healthy periods (n=54).

|  | All samples (n=117) | Infection samples (n=63) | Healthy samples (n=54) |
| --- | --- | --- | --- |
| **Maternal Age (years)** | | 21-40 (30.8 ± 4.6) | |
| **Parity** | | 1-3 (1.5 ± 0.7) | |
| **Fat content (%)** | 2.4-13.2 (6.4 ± 2.5) | 2.4-13.2 (6.1 ± 2.6) | 3.6-11.2 (6.9 ± 2.4) |
| **Milk cells/mL milk** | 111,111-3,517,241<br>(657,097 ± 621,040) | 111,111-3,517,241<br>(655,061 ± 799,468) | 200,000-1,305,556<br>(65,9472 ± 33,1935) |
| **Cell viability (%)** | 93.5-100 (98.2 ± 1.3) | 93.5-99.6 (97.9 ± 1.5) | 96-100 (98.4 ± 1.2) |
| **Total RNA (μg/mL)** | 0.000244-7.87 (0.58 ± 1.07) | 0.000244-2.81 (0.47 ± 0.74) | 0.000254-7.87 (0.68 ± 1.32) |
| **260/280 ratio** | 1.02-2.1 (1.81 ± 0.35) | 1.11-2.1 (1.81 ± 0.36) | 1.02-2.09 (1.82 ± 0.33) |
| **Known miRNA species\*** | 751-1,168 (1,017 ± 98) | 751-1,168 (1,022 ± 119) | 848-1,087 (1,005 ± 76) |
| **Novel miRNA species\*** | 16-167 (84 ± 28) | 16-110 (80 ± 27) | 66-167 (89 ± 28) |

\* The number of miRNA species was determined using deep sequencing in a subgroup of HM cell and fat samples (n=26).

### Processing of human milk and RNA extraction

Prior to milk fractionation for RNA/miRNA extraction, fat content of each whole milk sample was measured in a milk aliquot using the Creamatocrit method as previously described [44]. After this, whole milk samples were immediately fractionated as previously described [28,45], to obtain the three main milk fractions (cells, fat, and skim milk). Cells were counted using a Neubauer haemocytometer (BRAND, Wertheim, Germany) and cell viability was measured via Trypan Blue exclusion, as previously described [45]. RNA was then extracted from cells, fat, and skim milk immediately without cryopreservation using different extraction kits as previously optimized [46]. Therefore, RNA was extracted from milk cells (n=39) using the miRNeasy mini-Kit (Qiagen, Hilden, Germany), from milk fat (n=39) using the miRCURY RNA Isolation-Biofluids Kit (Exiqon, Vedbaek, Denmark), and from skim milk (n=39) using the miRCURY RNA Isolation- Cell&Plant Kit (Exiqon, Vedbaek, Denmark). NanoDrop 2000 Spectrophotometer (Thermo Scientific, Wilmington, MA, USA) was used to measure the concentration and purity of the total extracted RNAs. An Agilent Bioanalyzer 2100 instrument (Agilent, Santa Carla, CA, USA) with an RNA 6000 Nano Chip kit was used to measure the RNA concentration only for the samples used for small RNA sequencing. All miRNA samples were immediately stored at -80°C for small RNA sequencing and qRT-PCR.

### Multicolour Flow Cytometry

Multicolour flow cytometry was conducted on each HM cell fraction sample to examine the percentage contribution of leukocyte subsets to total milk cells through combined fluorescent-labelled antibodies against CD45-FITC, CD3-PE, CD4-APC, CD8-APC, CD56-APC, CD19-PE, and CD14-APC (BD Biosciences, San Diego, USA) (Miltenyi Biotec, Bergisch Gladbach, Germany) as described previously [6]. Briefly, freshly isolated and PBS-washed HM cells were further washed with 2% fetal bovine serum (FBS) (Gibco, NY, USA) in phosphate buffered saline (PBS, Gibco, Foster, CA), and were distributed into sterile 1.5-mL tubes. Fluorescent-labelled antibodies were added and incubated for 30 min at 4°C in the dark. An unstained control and respective isotype controls were used to account for background fluorescence. The cells were washed twice in 2% FBS in PBS, and were fixed in 1.5% paraformaldehyde/1% sucrose solution in PBS. Measurements were conducted with a FACS Calibur Flow Cytometer (Becton Dickinson, Franklin Lakes, NJ, USA). In depth analyses of leukocyte populations were conducted posthoc using FlowJo (http://www.flowjo.com).

### Small RNA sequencing and data analysis

Small RNA sequencing was conducted in a subgroup of HM cell and lipid samples (n=26), of which n=12 samples were collected when the mother and infant were healthy and n=14 samples during an infection episode. Skim milk samples were not sequenced due to the low amounts of RNA obtained. Sequencing libraries were prepared from HM cells (n=13) and fat (n=13) samples (both normal and abnormal) using the Solexa small RNAs protocol as previously described [29,47,48]. Briefly, 18-30 nucleotides long small RNA sequences were obtained by using polyacrylamide denture gels for size fractionation, and ligated to 5′-RNA and 3′-RNA adapters. qRT-PCR was used to reverse transcribe all the sequences into cDNA. cDNA was purified using PAGE gel for PCR amplification. After purification of the cDNA products, a SE49 lane was used on Illumina HiSeq2000 to sequence all small RNAs (Illumina, San Diego, CA, USA). Standard bioinformatics analysis including cleaning and filtering analyses was conducted on the raw tags/reads generated from the sequencer. Contaminated and low-quality reads were removed as previously described [28,29]. Then, all the resulted high quality small RNA reads were checked by determining each nucleotide base percentage/quality distribution on each nucleotide position. The clean reads were distributed based on the nucleotide size (18-30nt in length). All the clean reads were mapped to human genome using the SOAP software to analyse their expression and distribution using the latest version of human genome reference (hg38). All the clean and mapped reads were annotated using BLAST to Rfam and GenBank databases to determine and remove rRNA, tRNA, snRNA, scRNA, snoRNA, other ncRNAs, and degraded fragments of mRNAs. The remaining clean reads were matched using BLAST to the latest version of miRBase 21.0 (release August 2014) (http://www.mirbase.org/) to identify the human mature known miRNA species. The clean reads that mapped to the human genome but did not match to any known RNAs including miRNAs were used for novel miRNA prediction analysis. The Mirdeep software was used to structure the potential precursor miRNAs, the Dicer cleavage site, and the minimum free energy for all the nominated mature novel miRNAs. Finally, the nucleotide length on each position and base bias on the first position were determined for each read for all the nominated novel miRNA predicted.

### qRT-PCR validation

Six highly expressed mature known immune-related miRNAs were selected to validate their presence and expression level in all HM fractions (cells, fat, and skim milk) using qRT-PCR. TaqMan miRNA assays (Thermo Fisher Scientific, Waltham, MA, USA) were used to quantify let-7f-5p, miR-141-3p, miR-148a-3p, miR-181a5p, miR-182-5p, miR-375, and miR-99b-5p. Due to the low amount of RNA obtained from skim milk samples, all HM fractionated samples used for qRT-PCR experiments (n=85) obtained from 16 mothers were standardized to 70 ng/μL to accommodate for comparative analysis of skim milk miRNA together with cellular and fat miRNA. An endogenous miRNA (has-miR-30a-5p) was used as reference gene for analysis. Raw Ct values and relative quantitation (RQ) values were obtained using ABI 7500 software V2.0.6.

### miRNA differential expression analysis

The generated known and novel mature miRNAs obtained from sequencing were used for differential expression analysis between either the healthy and infection cohorts, or the milk cell and fat fractions. The normalisation formula: *normalised expression = actual miRNA count / total count of clean reads * 1,000,000* was used to obtain the expression levels of transcript per million (TPM). Fold change and p values were calculated using the following formulae: *Fold change = log_2_(normalised expressed miRNAs from a milk cohort / normalised expressed miRNAs from another milk cohort,* and p-value:

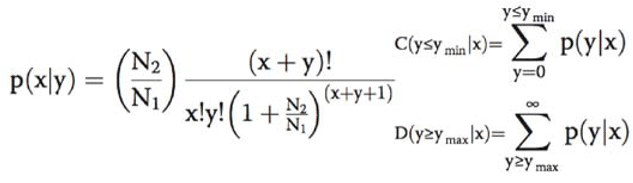

N1 represents the total reads in a given sample/cohort; x represents the total normalised reads of a given miRNA species; N2 represents the total reads of the given sample from another comparison cohort; and y represents the total normalised reads of a given miRNA species from another comparison cohort. DEGseq, an R package for identifying differentially expressed genes from RNA-seq data, was used to generate scatter plots and accurately determine p-values based on the above fold change formula [49]. p<0.05 of a given miRNA was considered as a differentially expressed value between two comparison cohorts.

### miRNA target prediction and functional analysis

Three different databases/algorithms were used to accurately predict the target genes of the top 50 most highly expressed miRNAs and top 50 differentially expressed miRNAs. These databases were targetscan (http://www.targetscan.org/), miRanda (http://www.microrna.org/microrna/home.do), and RNAhybrid (http://bibiserv.techfak.uni-bielefeld.de/rnahybrid). The prediction approach was based on the binding site or the seed region (6-8nt seed mir region) between a mature miRNA and a gene (mRNA). The identified target genes were also used for computational prediction functions and roles of these miRNAs/targets. Two databases were used to determine miRNA functions, Gene Ontology (GO) (http://www.geneontology.org/) [50] and the Kyoto Encyclopedia of Genes and Genomes (KEGG) (http://www.genome.jp/kegg/) [51]. GO was used to predict roles of different signaling and cellular function ontologies (cellular component, molecular function and biological process), while KEGG was used for prediction of metabolic and cellular pathways.

### Statistical analysis

The statistical analysis of the RQ values obtained using qRT-PCR was done using R Studio Version 0.98.1103 package [52] between the comparison cohorts using linear mixed effects (LME) models, where p≤0.05 was considered statistically significant. All participants were grouped in three groups according to their health status for total cell number, leukocyte number, and miRNA concentration analysis between and within HM fractions. These groups were HM samples obtained during healthy conditions (n=17 and during infections including common cold/flu (n=10), mastitis (n=3), cracked nipple, (n=1), toe infection (n=1), gastrointestinal infection (n=3), recurring ear infection (n=3), and a mother suffering from ulcerative colitis/sweets syndrome/Hashimoto’s disease (n=2) (total of infection samples: n=23). R Studio Version 0.98.1103 package was used to perform the statistical analysis via general linear hypothesis tests, and linear mixed effects modeling (LME) via nlme [53] and lattice [54] packages. If p<0.05, the comparison between two parameters was considered significantly different. In addition, the difference of the number of identified miRNA species between different health cohorts or cell and fat samples was statistically tested using LME models to account for the possibility of individual variation. The verification with LME modelling indicated that there was no random effect due to individual variation in this study. Thus, linear regression modelling was used for the final comparison analysis.

### Availability of supporting data

All raw, analysed small RNA sequences were submitted to the NCBI Gene Expression Omnibus database under accession number GSE80546. Additional information is also included as supplementary files.

## RESULTS

### Associations between human milk cell, fat and RNA contents in relation to the health status of the mother/infant dyad

Total HM cells, miRNA concentration, and fat content obtained from all samples (n=117 from all HM fractions and both healthy and infected dayds) were tested for statistically significant associations in relation to health status **(Table 1)**. The RNA content did not differ between the common cold, other infections and healthy cohorts (p>0.05). No difference was found in total cell number per mL of HM between the healthy cohort and the cohort suffering from common cold (p=0.380) or other infections considered all together (p=0.07), or between the cohort with common cold and the other infections cohort (p=0.051). However, HM fat content was lower in the common cold compared to the healthy cohort (p=0.045), but it did not differ between the healthy and the other infections cohort (p=0.789). Further, RNA content (per million of cells or per μL of fat or skim milk) measured by NanoDrop was extremely higher in HM cells compared to the fat and skim milk fractions (p<0.001). Similarly, the HM fat fraction had higher RNA content than skim milk (p=0.001). A significant positive correlation was observed between HM fat content and RNA concentration of fat samples in the healthy cohort (p=0.011), but this was not seen in the common cold and other infections cohorts (p>0.05). Moreover, HM cell content was positively associated with HM fat content (p=0.034). Cellular RNA content was not related to the number of cells per mL of HM (p>0.05). Further, no difference was observed in cellular RNA content between the healthy and common cold cohorts (p=0.96), or in fat RNA content between the healthy and common cold cohorts (p=0.10), or in skim milk RNA content between the healthy and common cold cohorts (p=0.10).

### Infection of the mother/infant dyad stimulates a rapid response in leukocytes and leukocyte subsets of human milk

The HM cell content was positively associated with the number of leukocytes (% of total cells) (p=0.013) in both the common cold and other infection types. The relative number of HM leukocytes (% of total cells) was higher in the common cold cohort compared to the healthy cohort (p=0.023) **(Tables 2 and 3)**.

**Table 2.**
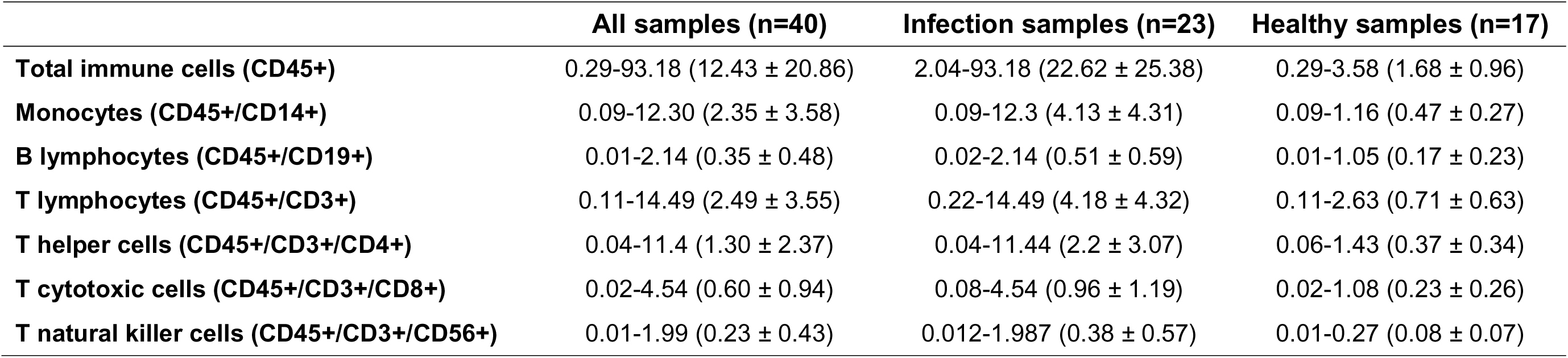
The range (mean percentage ± standard deviation, S.D.) of the total leukocyte count (% of total HM cells) and individual leukocyte subtypes counts (% of total HM cells) measured by flow cytometry (n=18 participants).

**Table 3.** The relationship between the number of the identified known and novel miRNA species and total HM cells, total leukocytes, and leukocyte subsets measured by flow cytometry.

| Relationship with known miRNAs | Healthy cohort<br>(n=48) | Common cold<br>cohort (n=30) | Other infections<br>cohort (n=39) |
| --- | --- | --- | --- |
| Total cell content (per mL of milk) | 0.779 | 0.291 | 0.332 |
| Total leukocytes (CD45+) | 0.555 | 0.091 | 0.240 |
| Monocytes (CD45+/CD14+) | 0.360 | **0.059** | 0.328 |
| B lymphocytes (CD45+/CD19+) | 0.721 | 0.993 | **0.027** |
| T lymphocytes (CD45+/CD3+) | 0.878 | 0.733 | **0.039** |
| T helper cells (CD45+/CD3+/CD4+) | 0.859 | 0.634 | 0.148 |
| T cytotoxic cells (CD45+/CD3+/CD8+) | 0.895 | 0.729 | 0.579 |
| T natural killer cells (CD45+/CD3+/CD56+) | 0.993 | 0.554 | 0.623 |
| Relationship with novel miRNAs | Healthy cohort<br>(n=48) | Common cold<br>cohort (n=30) | Other infections<br>cohort (n=39) |
| Total cell content (per mL of milk) | 0.576 | 0.240 | 0.525 |
| Total leukocytes (CD45+) | 0.777 | 0.079 | 0.616 |
| Monocytes (CD45+/CD14+) | 0.501 | **0.036** | 0.528 |
| B lymphocytes (CD45+/CD19+) | 0.725 | 0.925 | 0.883 |
| T lymphocytes (CD45+/CD3+) | 0.949 | 0.672 | 0.895 |
| T helper cells (CD45+/CD3+/CD4+) | 0.890 | 0.561 | 0.708 |
| T cytotoxic cells (CD45+/CD3+/CD8+) | 0.786 | 0.677 | 0.565 |
| T natural killer cells (CD45+/CD3+/CD56+) | 0.952 | 0.523 | 0.521 |

### Effect of maternal/infant infection on the number of miRNA species in human milk

Next generation sequencing (small RNA SE50 using Illumina HiSeq 2000) was used to profile miRNAs in a subgroup of HM cell and fat samples (n=26 of which n=13 cell and n=13 fat samples) from healthy and infected mother/infant dyads. Raw reads data from all samples were analysed to rid of all the contaminated reads, where 770,374,554 raw reads were generated from sequencer platform **(Figure 1A)**. Filter analysis was done to remove 81,091,772 (10.5%) reads, and 689,282,782 clean reads (89.5%) were retained to be used for the subsequent bioinformatics analysis **(Figure 1A- 1B)**. All clean small RNA reads were distributed based on their nucleotide base length, in which the vast majority of the clean reads (82.4%) were classified between 20 and 24 nucleotides sequence length **(Figure 1C)**. In particular, most of the clean reads (34.2%) had 22 nucleotides, which is typically the length of most miRNAs in mammals **(Figure 1C)**. Thereafter, all clean reads were also mapped to different RNA categories using short oligonucleotide alignment Package (SOAP) **(Figure 1D-1E)**. Clean small RNA reads were mapped to Genebank and Rfam to determine and remove tRNA, rRNA, snRNA, snoRNA, scRNA, repeat small RNAs, and sense/antisense of introns and exons **(Figure 1D-1E)**. The remaining total clean reads (589,903,296; 85.6%) were used to identify human mature known miRNAs by mapping to miRBase 21.0 database using BLAST, and to predict novel miRNAs using mirdeep software and other criteria **(Figure 1D-1E)**.

**Figure 1.**
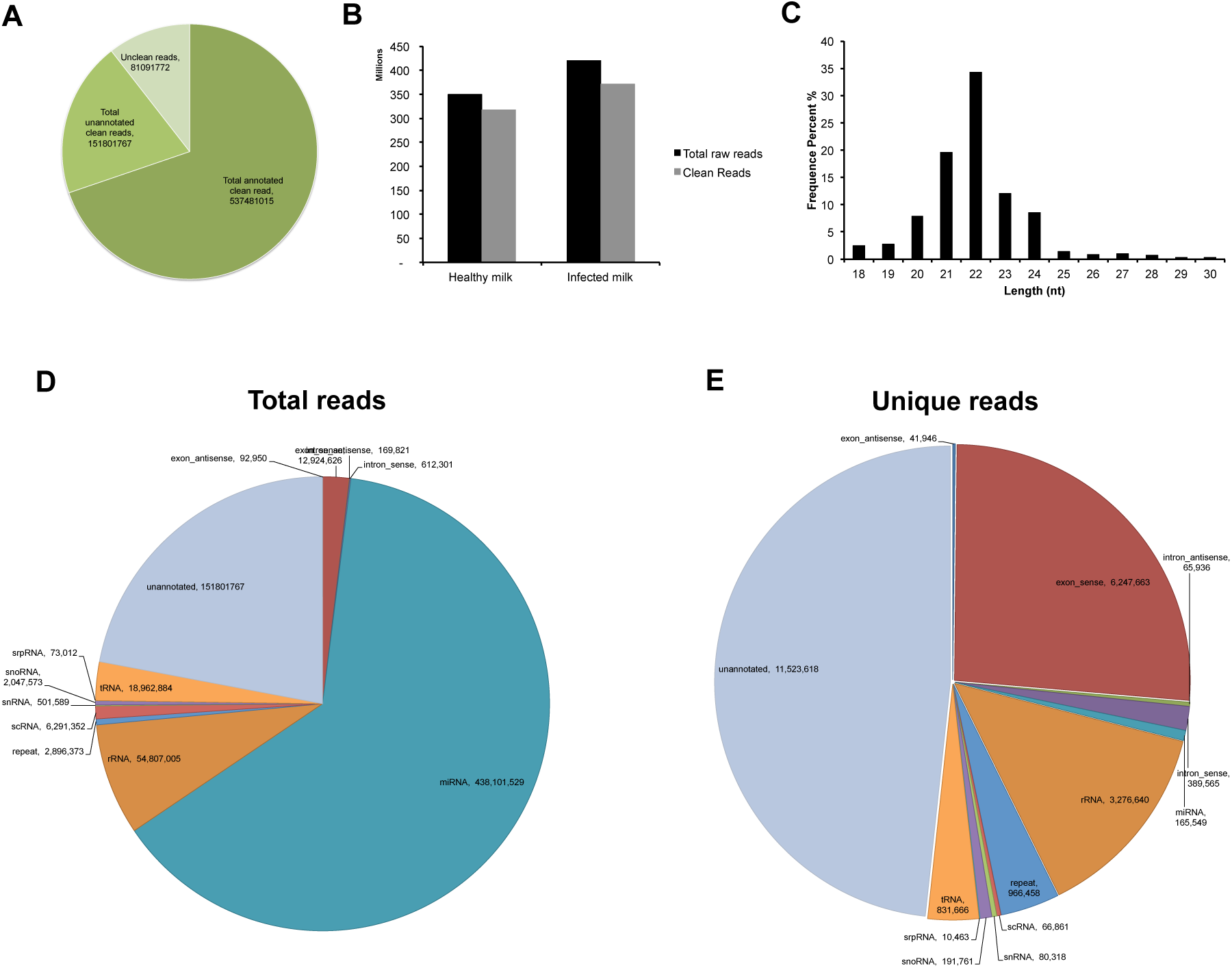
Filter and annotation of small RNA reads generated by Next Generation Sequencing. **(A)** All the generated small RNA reads statistics. **(B)** The number of raw reads compared to clean reads after filtering contaminated reads. **(C)** The distribution of small RNA clean reads based on their nucleotide length between 18 and 30 nucleotides. **(D)** Total small RNA clean reads were annotated to one or more RNA categories (i.e. rRNA, tRNA, scRNA etc.). **(E)** Small RNAs were mapped to only one RNA category using the following priority rule: rRNAetc (in which Genbank > Rfam) > known microRNA > piRNA > repeat > exon > intron3. rRNAs were used as a marker of sample quality, with the criterion of high quality when <40% in each sample.

The total matched reads to miRBase were 441,356,898 reads for all samples, where the average of the total matched reads per sample was ∼17M **(Supplementary Table S1)**. Thus, 1,780 mature known miRNAs were identified in HM cell and fat samples derived from healthy and infection cohorts. In particular, 1,680 known miRNAs were determined in the infection cohort, whilst 1,606 in the healthy cohort **(Supplementary Table S1)**. The total number of the profiled known miRNA species was not statistically different between the common cold and healthy cohorts (p=0.677) **(Table 1)**. The majority of the identified known miRNA species were common between the cohorts (1,506 common known miRNA species) **(Figure 2A)**. However, some known miRNAs with low expression were found to be specific to either the healthy or the infection cohort, with 100 known miRNAs (210 total reads) and 174 known miRNAs (467 total reads) being specific to the healthy and infection cohorts, respectively **(Figure 2A)**.

**Figure 2.**
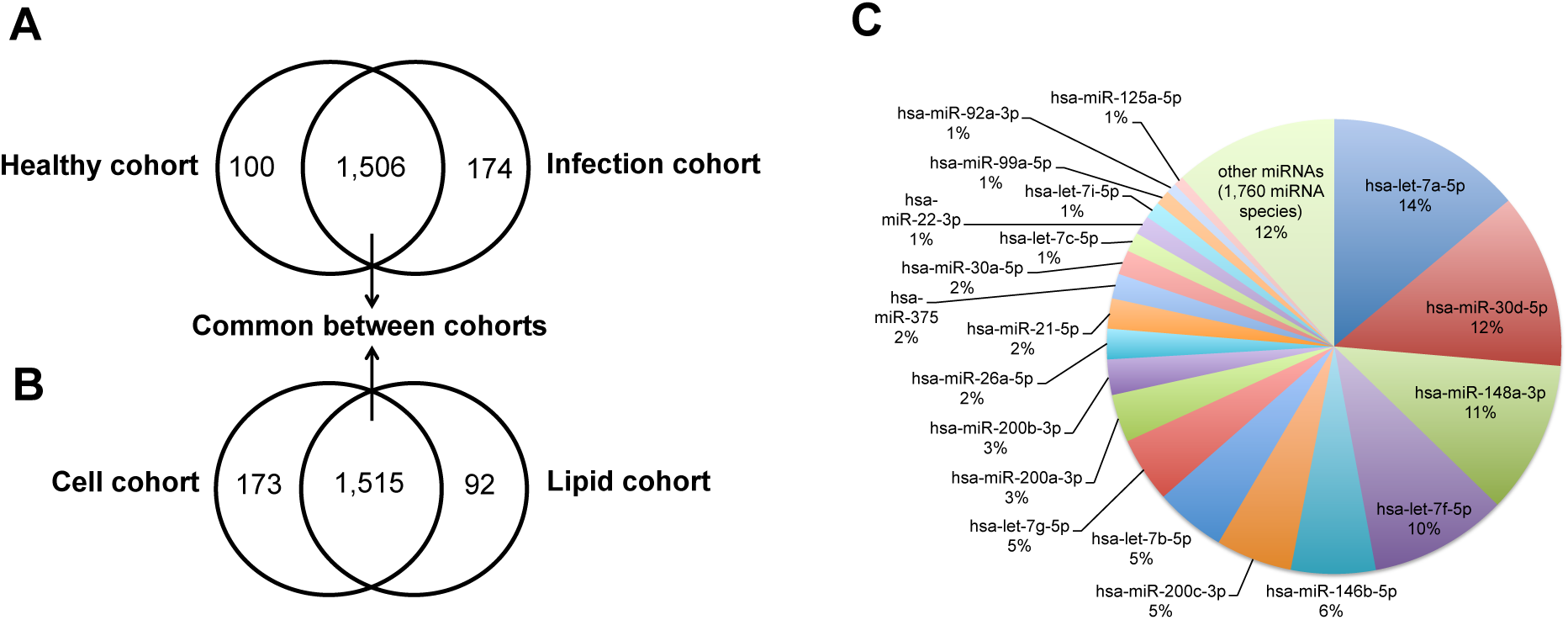
Characteristics of the identified miRNA species number between **(A)** healthy and infection cohorts, and between **(B)** cell and fat cohorts. **(C)** Pie chart showing the 20 most highly expressed miRNAs in all sequenced samples (n=26) compared to identified 1,760 miRNA species.

When the HM cell and fat samples were analysed separately, it was shown that the profiled miRNA species number in HM cell samples (n=13) was slightly greater than that in the HM fat samples (n=13), but this was not statistically significant (p=0.922). Specifically, 1,688 known miRNA species (range: 751-1,168) were identified in HM cells (total reads: 219,942,145; average reads per sample: 16,918,626), whilst 1,607 (range: 923-1,099) known miRNAs were profiled in HM fat (total reads: 221,414,753; average reads per sample: 17,031,904). Similar to the comparison between the healthy and infection cohorts, the majority of known miRNA species (1,515) were commonly determined in both cell and fat fractions of HM. However, 173 and 92 known mature miRNAs with low expression were specific to the HM cell and fat fractions, respectively **(Figure 2B)**. The total RNA content (ng/1 million of cells) of HM cells measured by NanoDrop was positively associated with the number of known miRNA species obtained in cells in the healthy cohort (p=0.027), but not in the common cold or other infection cohorts (all p>0.05). HM fat content was not related to the number of known miRNAs in any of the cohorts (healthy or infection) (all p>0.05).

The top 20 known mature miRNAs out of 1,780 determined in both the cell and fat fractions of HM in the healthy and infection cohorts contributed 88.3% of all the identified known miRNAs **(Figure 2C; Supplementary Table S1)**. The most highly expressed miRNA in all samples (n=26) was hsa-let-7a- 5p with total reads 61,429,206, contributing 13.9% of all the identified known miRNAs in all sequenced samples (n=26) **(Figure 2C; Supplementary Table S1)**. Six members of the let-7 family (let-7a-5p/7f-5p/7b-5p/7g-5p/7c-5p/7i-5p) were in the top 20 known miRNAs, and let-7d-5p and let-7e-5p were also ranked in the top 50 miRNAs **(Supplementary Table S1)**.

For the discovery of novel miRNA species, small RNA reads obtained from all HM cell and fat samples (n=26) that unannotated to any RNA categories and mapped to the human genome were used to predict novel miRNAs **(Supplementary Figure S1; Supplementary Table S2)**. Therefore, 151,801,767 (22% of the total clean/filtered reads) total clean and unannotated reads were nominated as novel miRNAs, and examined with different prediction criteria **(Figure 1A)**. First, nucleotide base bias on the first position with certain length and on each position was assessed for all total unannotated small RNA reads **(Supplementary Figure S2A-S2B)**. The second step was to structure the stem loop (precursor miRNA) of the predicted mature miRNA sequences using mirdeep, and to determine the Dicer cleavage site and the minimum free energy score (mfe). In all samples (n=26), less than 0.05% of the total unannotated reads (151,801,767) matched the above strict criteria for novel miRNA prediction. In detail, 592 novel mature miRNA sequences were predicted, with only 65,878 total reads **(Supplementary Figure S2)**. Amongst the total reads of the novel miRNAs, the top 20 novel miRNAs were found to contribute 73.3% (total reads 48,295) of the total reads (65,878) **(Table 4)**. In addition to the above criteria, high-confidence novel miRNAs were determined in order to obtain highly predictable novel miRNAs amongst the total number of 592 predicted novel miRNAs. miRNAs of high confidence were identified in ≥3 samples (n=26) with total reads ≥20. This yielded 95 novel mature miRNAs of high confidence **(Supplementary Figure S2)**. Among all determined novel miRNAs, 122 unannotated miRNA sequences were 22 nucleotides in length, whilst 54 and 118 novel miRNA species were 23 and 21 nucleotides long, respectively **(Supplementary Figure S2)**. Further, precursor miRNAs (pre-miRNAs) were structured for all the discovered novel mature miRNAs using mirdeep software. The top 20 highly expressed miRNAs are shown in **Supplementary Figure S3**.

**Table 4.**
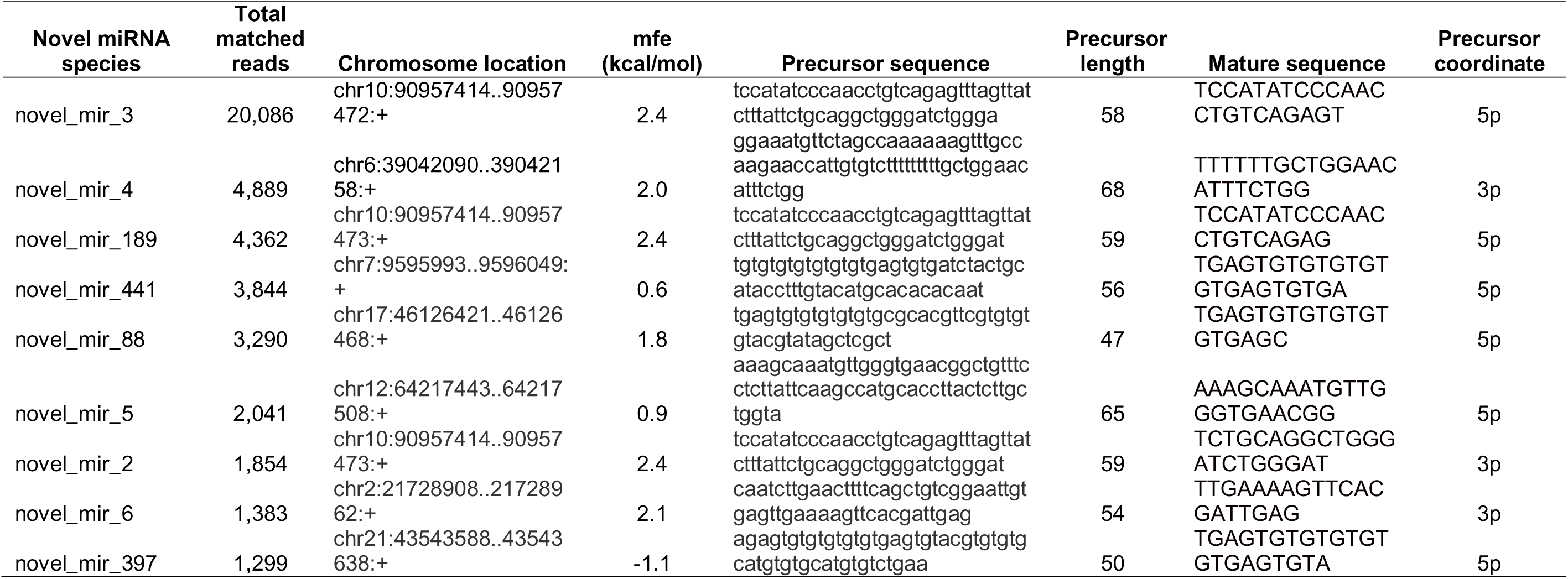
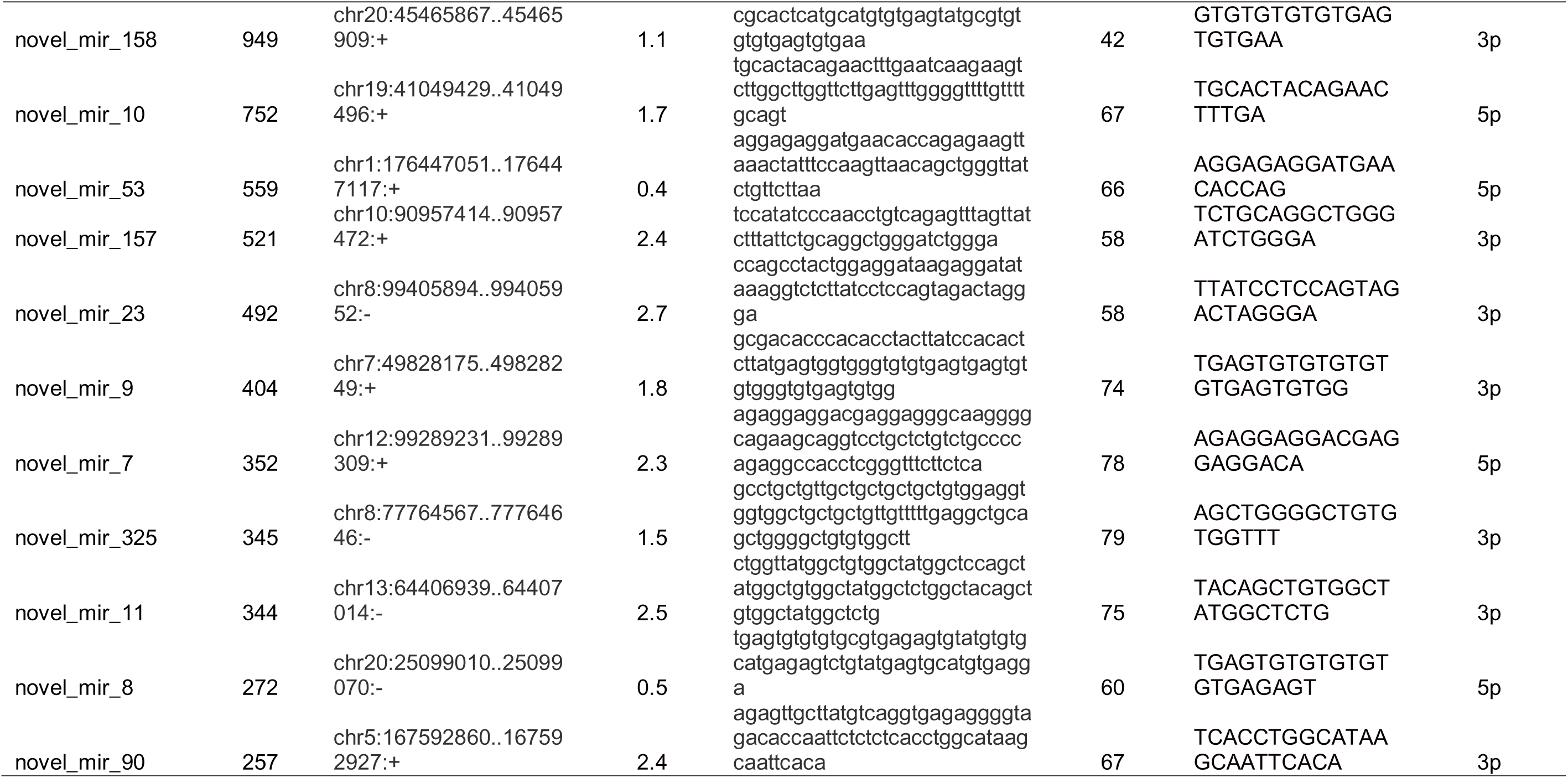
The top 20 highly expressed novel mature miRNA species in all sequenced samples (n=26), with their total reads and locations mapped to the human genome. Predictable precursor and mature sequences and their minimum free energy (mfe) score are included for each of the top 20 novel miRNAs.

Statistical comparisons showed similar total number of novel miRNA species between HM cells and fats (p=0.36), and between the healthy and the common cold cohorts (p=0.419). The number of novel miRNA species obtained from the healthy, common cold and other infections cohorts was not related to HM RNA cell content (p>0.05) or HM fat content (p>0.05).

### Human milk miRNAs are differentially expressed in response to infections of the mother and/or infant during breastfeeding

Fold change/Log2Ratio was calculated from the normalised miRNA expression level obtained from two cohorts to identify the degree of difference, which was used to rank the most different miRNAs, where -2≤Log2Ratio≥2. The DEGseq R statistical package was used to determine whether the miRNA expression was significantly different between two cohorts. The expression of 453 known miRNAs (25.5% of all known miRNAs sequenced) was significantly different (p<0.05) between healthy (n=12) and all infection cohorts (n=14) **(Figure 3A; Supplementary Table S3)**. Of these, 70 known miRNAs had significant fold change values (-2≥fold change≤2), with 62 known miRNAs upregulated and 8 known miRNAs downregulated in the all infections cohort compared to the healthy cohort **(Supplementary Table S3)**. When HM cells and fat were analysed separately, 447 and 334 known miRNAs were differentially expressed (p<0.05) between the healthy and infections cohorts of cells and fats, respectively **(Figure 3B-3C; Supplementary Table S4-S5)**. Of these, 66 miRNAs were highly different (-2≥fold change≤2) between HM cell samples from the infections cohort (n=7) and HM cell samples from the healthy cohort (n=6), with 62 known miRNAs (93.9%) upregulated in infection HM cell samples. Similarly, 53 known miRNAs were highly different between HM fat samples from the infection cohort (n=7) and the healthy cohort (n=6), with 41 known miRNAs (77.4%) being upregulated in infection HM fat samples **(Supplementary Table S4-S5)**. In a comparison between infection cell samples (n=7) and infection fat samples (n=7), it was revealed that HM cells had more upregulated miRNAs in response to infection (32 miRNAs - 88.9%) compared to HM fat, which conserved only 4 upregulated miRNAs **(Figure 3D; Supplementary Table S6)**. On the other hand, 389 miRNAs were differentially expressed between HM cells and fat from the healthy cohort. Of these, 57 miRNAs were considered as highly different miRNAs, with 48 miRNAs (84.2%) being upregulated in HM cells compared to HM fat from the healthy cohort **(Figure 3E; Supplementary Table S7)**.

**Figure 3.**
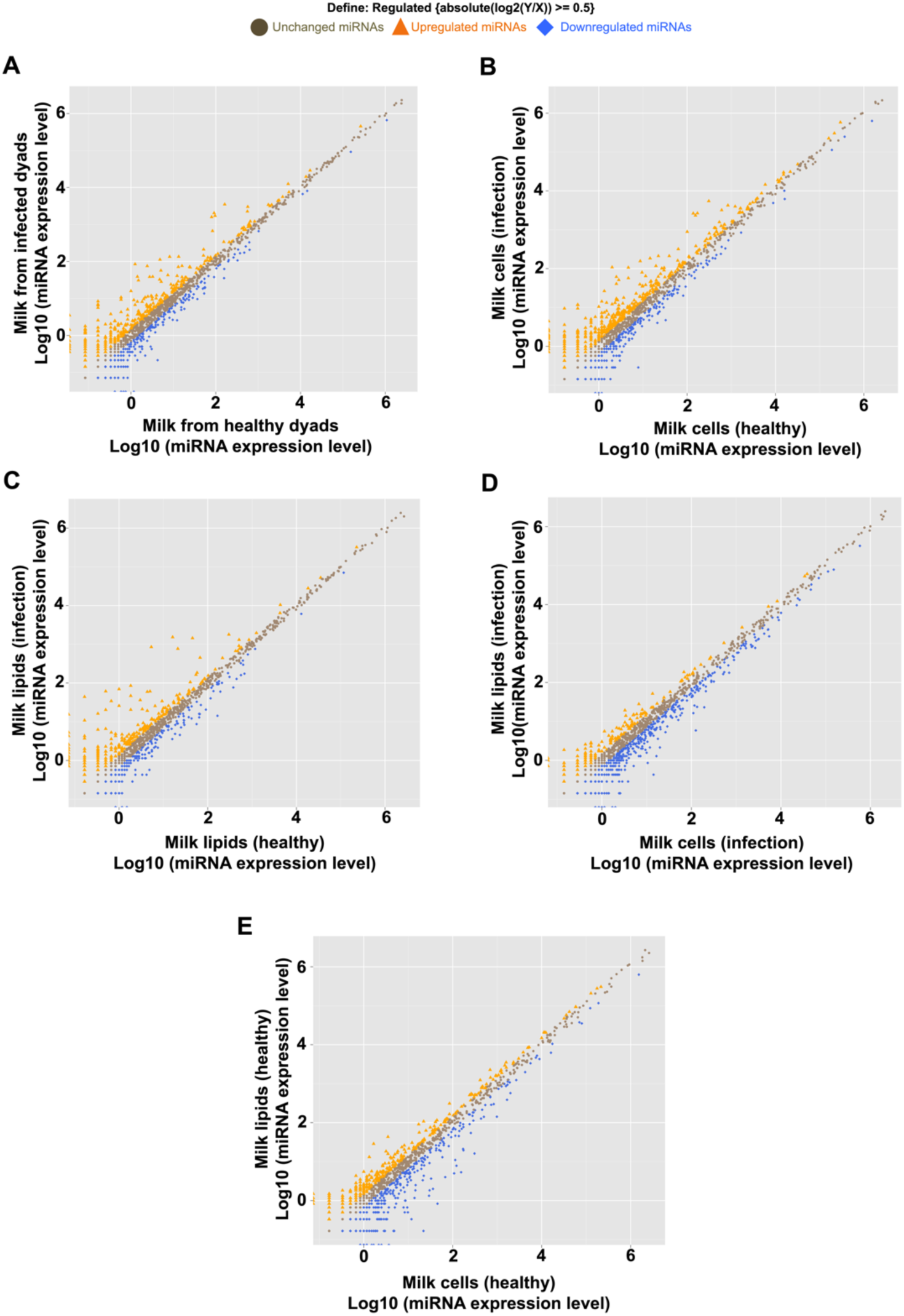
Scatter plots showing the differentially expressed human milk known miRNAs between healthy and infection sample cohorts. **(A)** The differentially expressed miRNAs between all healthy/cell and healthy/fat samples (n=12) versus all infection/cell and infection/fat samples (n=14), between **(B)** healthy/cells (n=6) and infection/cells (n=7), between **(C)** healthy/cells (n=6) and healthy/fat (n=6), between **(D)** infection/cells (n=7) and infection/fat (n=7), and between **(E)** healthy/fat (n=6) and infection/fat (n=7).

Similarly to known miRNA species, differences in expression with infection were also seen for the novel miRNA species. Specifically, 268 novel miRNAs (48.3% of total novel predicted miRNAs) were significantly differentially expressed between the healthy cohort (n=12) and the infection cohort (n=14) **(Supplementary Figure S4A; Supplementary Table S8)**. Among these, 258 novel miRNAs were highly different between the two cohorts, and the majority of them where upregulated in the infection cohort (163 miRNAs; 63.2%) compared to the healthy cohort **(Supplementary Table S8)**. In addition, 158 novel miRNAs were differentially expressed between HM cells (n=7) and HM fat (n=7) from the infection cohort **(Supplementary Figure S4B; Supplementary Table S9)**. Of these, 140 novel miRNAs were found with high degree of difference, with almost half of them being upregulated in HM cells from the infection cohort (71 miRNAs; 50.7%) **(Supplementary Table S9)**. Comparisons between HM cells (n=6) and fat (n=6) from the healthy cohort yielded 225 differentially expressed novel miRNAs **(Supplementary Figure S4C; Supplementary Table S10)**. Most of these were upregulated (131 miRNAs; 58.2%) in HM cells compared to fat from the healthy cohort **(Supplementary Table S10)**.

Heat map analysis showed that most of the top 50 known miRNAs were similarly expressed between the healthy and infection cohorts, with some of these miRNAs being differentially expressed (p<0.05). Hierarchical clustering analysis revealed higher correlations within cell samples and within fat samples **(Figure 4A)**. In contrast, most of the top 50 novel miRNAs were differentially expressed between the healthy and infection cohorts due to most of these being expressed only in few samples, i.e. being specific to the mother/infant dyad **(Figure 4B)**. A moderate correlation was seen between the HM cell and fat fractions **(Figure 4A)**. In regards to the top 50 differentially expressed known miRNAs, the vast majority of them were upregulated in the infection cohort compared to the healthy cohort **(Figure 4C-4E)**. Further, most of these were upregulated in the HM cell fraction compared to the fat fraction **(Figure 4F- 4G)**.

**Figure 4.**
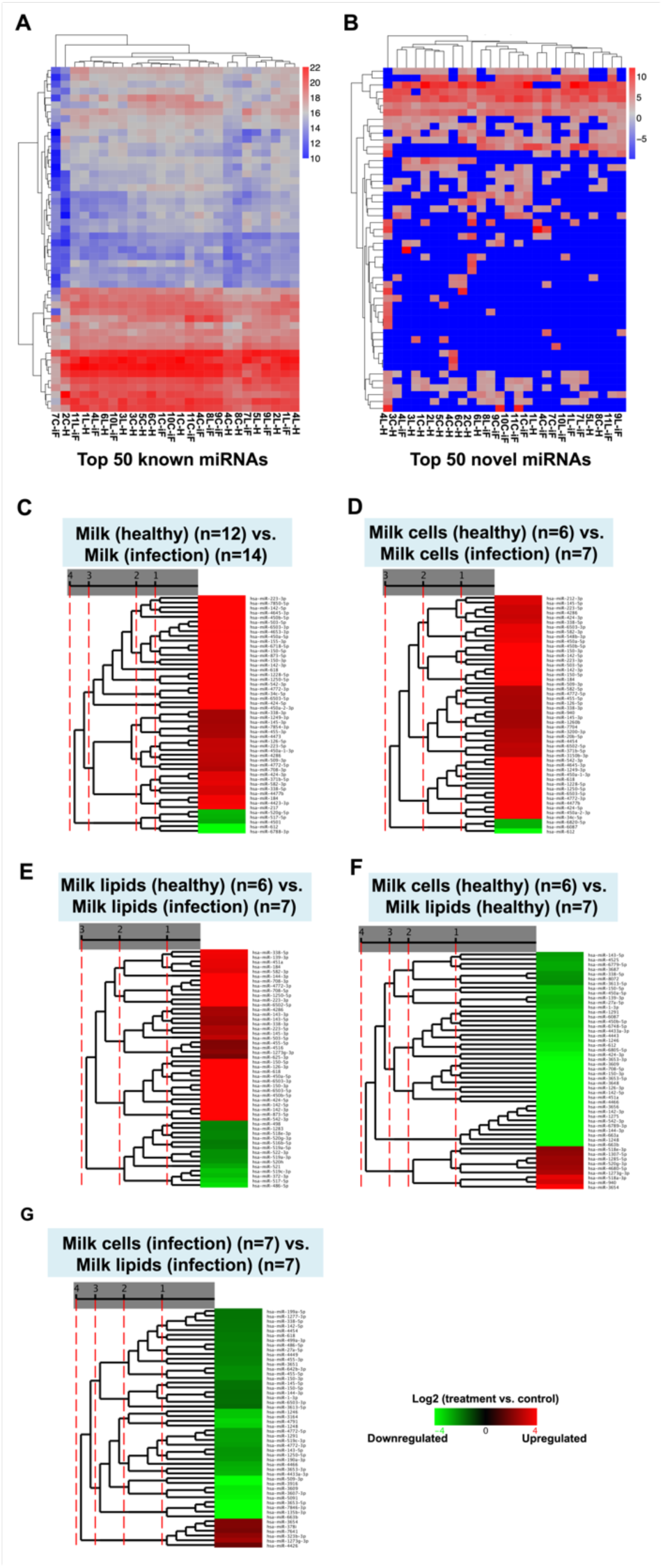
Hierarchical clustering linkage of the **(A-B)** top 50 most highly expressed known and novel miRNAs in all human milk cell and fat samples (n=26). **(C-G)** Clustering of the top 50 differentially expressed known miRNAs between healthy and infection samples (red indicates the upregulated miRNAs between two sample cohorts).

### Associations between human milk leukocytes and miRNAs during healthy and infection periods

**Table 4** shows the statistically explored associations between the number of known or novel miRNA species and the percentage of leukocytes and leukocyte subsets for the healthy, common cold and other infections cohorts. Although for the majority of these associations, no significance was observed, the number of known miRNA species in HM cells of the other infection cohort (which did not include the common cold) was positively related to the percentage of B and T lymphocytes out of total cells (p=0.027 and p=0.039, respectively) **(Table 4)**. A positive association was also found between the number of known and novel miRNA species and the percentage of monocytes (p=0.059 and p=0.036, respectively) **(Table 4)**.

### Validation of highly expressed miRNAs in different human milk fractions during healthy and infection periods

Six mature known miRNAs (let-7f-5p; miR-141-3p; miR-148a-3p; miR-181a5p; miR- 182-5p; miR-375; miR-99b-5p) were selected for qRT-PCR validation in all three fractions of HM, including the cells, fat and skim milk. These miRNAs are known to play roles in immune response to different bacterial and viral infections, and to participate in the development of the immune system **(Supplementary Table S11)**. All these miRNAs were identified in all the samples and different HM fractions (n=67) examined, where HM cells showed mostly higher expression levels compared to HM fat and skim milk, with the fat fraction being higher than the skim milk, though not statistically significant (p>0.05) **(Figure 5)**. Although infections influenced to some extent the levels of expression of all six miRNAs tested, no statistically significant difference was observed between the healthy and infection cohorts (p>0.05) **(Figure 5)**.

**Figure 5.**
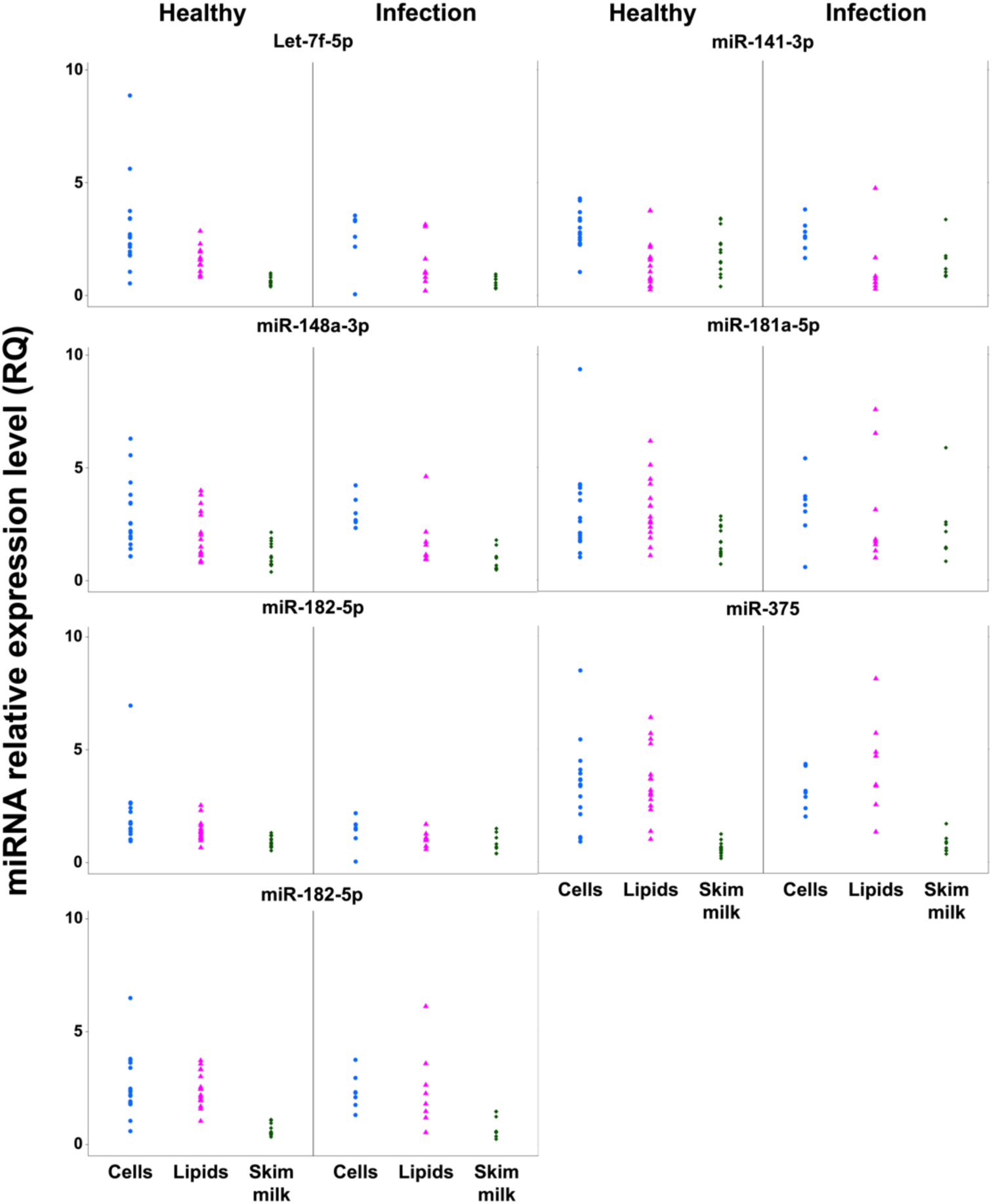
The expression patterns of 7 highly expressed immune-related known miRNAs in the three human milk fractions (cells, fat and skim milk) and between healthy and infection cohorts using qRT-PCR. The Y axis indicates the relative expression (RQ) of the miRNAs to the endogenous control (miR-30a-5p), whilst the X axis indicates the milk fractions in healthy and infection samples. 67 milk samples obtained from 16 mothers were used for qRT-PCR validation. Linear mixed effects modeling revealed no significant differences (p>0.05) in known miRNA expression patterns between different milk fractions and between healthy and infection cohorts.

Moreover, mastitis influenced expression of certain miRNAs, although the sample number was not enough to examine statistical significance of this effect **(Figure 6A- 6B)**. Except miR-182-5p in the HM fat fraction, all other miRNAs in both cell and fat fractions (n=12) showed different expression levels between the three stages of mastitis sampled (from the initiation of the infection to the recovery period) and the healthy sample from the same participant. Let-7f-5p was expressed at extremely low and high levels in mastitis compared to healthy status in HM cells and fats, respectively **(Figure 6A-6B)**. Further, a HM sample obtained when the participant experienced cracked nipples, and which was associated with increased number of leukocytes compared to HM collected during a healthy period from this participant **(Figure 6C)**, showed large differences in expression of the majority of tested miRNAs, with the exception of miR-182-5p **(Figure 6C)**. Further, PCA analysis was conducted to explore the relationship of the expression patterns of the 7 miRNAs tested between the different HM fractions (cells, fat, and skim milk). This demonstrated that most of the samples from each HM fraction are clustered together **(Figure 6D)**.

**Figure 6.**
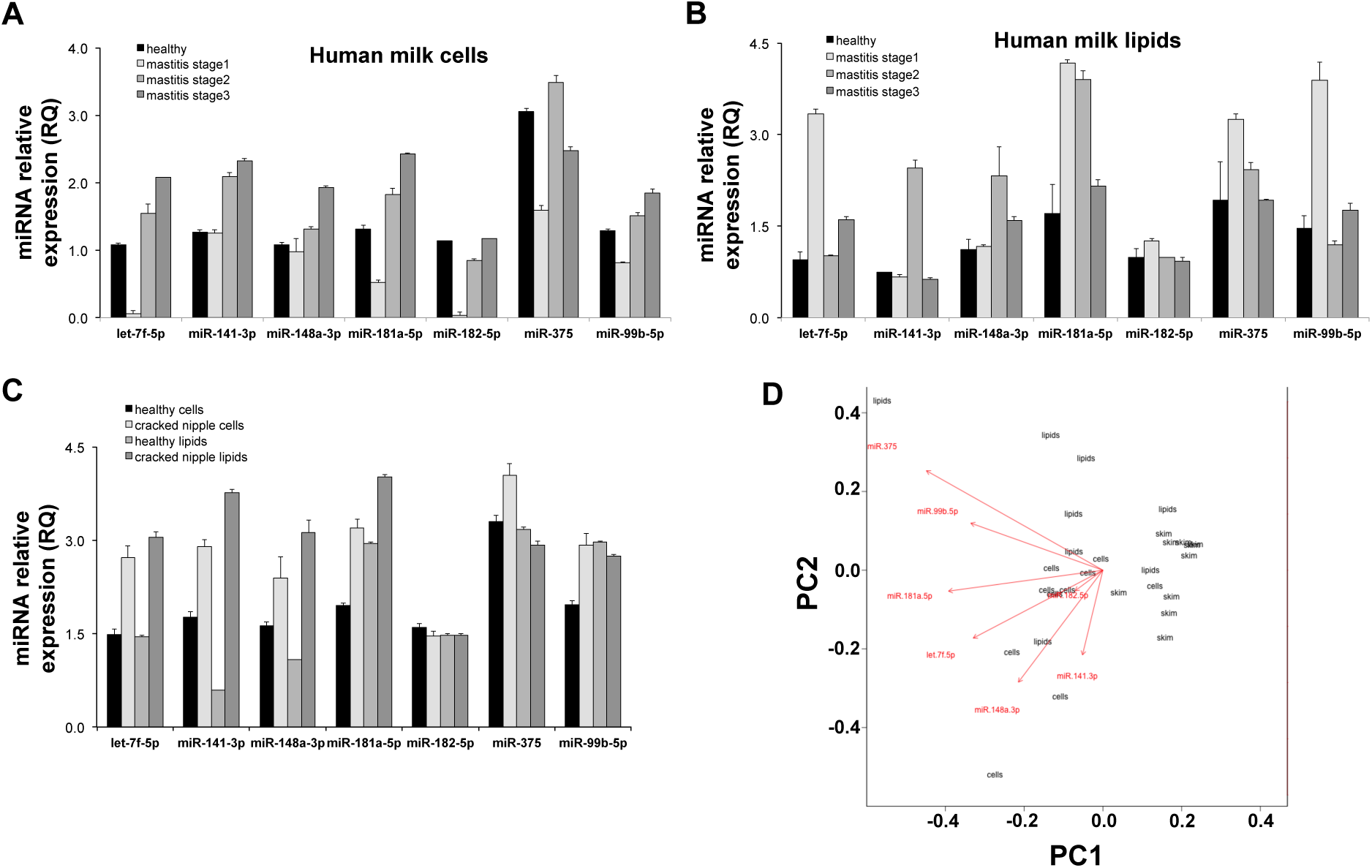
Bar charts showing the expression level of 7 highly expressed immune-related known miRNAs between healthy and mastitis samples (n=12) obtained from the same mother in **(A)** human milk cells and in **(B)** human milk fat. **(C)** Expression patterns of the same 7 miRNAs between healthy and cracked nipple samples (n=6) obtained from the same mother. **(D)** PCA analysis indicating the correlation between the three human milk fractions in expression levels measured by qRT-PCR.

### Prediction of miRNA target genes

The majority of the protein coding genes available in NCBI was found to be targeted by the top 50 most highly expressed and the differentially expressed known miRNAs between healthy and infection cohorts. RNAhybrid, TargetScan and miRanda software/tools were used to predict the gene targets for the assigned miRNAs. The top 50 known miRNAs in all samples (n=26) targeted 36,308 unique mRNAs **(Supplementary Table S12A)**. 36,224 genes were predicted to be the targets of the top 50 differentially expressed known miRNAs **(Supplementary Table S12B)**. In regards to the novel miRNAs, most of the top 50 and differentially expressed novel miRNAs were homologous. Therefore, the gene targets were identical for the entire top 50 highly expressed and the top 50 differentially expressed novel miRNAs, with 36,265 genes predicted for all of these novel miRNAs **(Supplementary Table S13)**.

GO and KEGG were used to investigate the predictable functions and roles of the top 50 differentially expressed known miRNA targets **(Supplementary Table S14- S15)**. 3,235 immune system processes related genes were found to be targeted by these differentially expressed miRNAs **(Supplementary Table S14)**. Further, targets were determined as biological regulation, growth and developmental processes-related miRNAs, including immune system and immune response in these mechanisms **(Supplementary Table S14)**. KEGG was used to identify the metabolic processes associated with these differentially expressed miRNAs. miRNA gene targets were shown to be involved in many immune responses from bacterial to viral infections **(Supplementary Table S15)**. Of these, 221 targets were involved in the pathway of response to *Staphylococcus aureus* infection. 104 targets of the top 50 differentially expressed miRNAs were involved in intestinal immune network for the IgA production pathway **(Supplementary Table S15)**.

## DISCUSSION

In the last decades, increasing evidence has demonstrated unique attributes of human milk (HM), shifting the dogma considering it a mere medium of nutrition into a live biofluid rich in regulatory signals, both cellular and molecular [22,45,55]. A plethora of HM components have been associated with provision of protection as well as developmental programming. Immunomodulatory agents are abundant in HM and include both molecules such as immunoglobulins, lactoferrin, and lysozyme, and immune cells ^10^. Leukocytes have been primarily examined in human colostrum as well as in milk of other mammalian species, such as the dairy cow, however it is only recently that more advanced single cell analytical methods have been employed to shed light into the leukocytic content of HM and factors influencing it [10]. Studies based on flow cytometric analysis of milk leukocytes and their subsets have demonstrated that although human colostrum is dominated by leukocytes in accordance with the increased protection needed by the newborn, mature HM is rich in epithelial cell signatures, with leukocytes contributing up to ∼2.5% of total milk cells when both the mother and infant are healthy [6,13,56,57]. Instead, during periods of infection of either the mother or the infant, HM leukocytic content significantly and rapidly increases to facilitate recovery and provide active immunity to the infant [6,13]. Therefore, it has been proposed that the leukocyte content of HM can be used as an indicator of the health status of the lactating mammary gland as well as the breastfeeding mother and infant [6,10]. Similarly, novel molecules called miRNAs recently discovered in mammalian milk have been suggested to be of diagnostic value and provide immunological signals associated with the health status of the mammary gland as well as the infant [22,33,43,58,59]. Milk miRNAs have been shown to be highly resistant under harsh conditions, and have been thus hypothesised to survive the infant’s gastrointestinal tract and enter the bloodstream. At the same time, both human and animal studies have examined milk miRNAs [14,20,27–30,32,34,46,60], revealing various factors, such as the stage of lactation and milk removal during breastfeeding, that can influence the miRNA content and composition of HM, hence suggesting pleiotropic functions for these molecules in the mother’s gland and in the infant [27–29,46]. Outside the lactation field, miRNAs have been used as biomarkers for many normal and abnormal conditions, such as cancers and immune disorders, due to their high sensitivity in response to any changes occurring in human body at both the cellular and molecular levels [26,61,62]. A range of miRNAs such as mir-155 and let-7 have been demonstrated as essential modulators of immunity and immune response during infection, including development and differentiation of B and T cells (adaptive immune response), and proliferation of monocytes and neutrophils (innate immune response) [26,63,64]. Although studies in the dairy cow have demonstrated responses of milk miRNAs to infection of the udder (mastitis) [33–35], investigations of the potential immunomodulatory functions of HM miRNAs in response to infection of the mother/infant dyad are lacking. To address this, we examined the miRNA profile of different fractions of HM (cells, fat and skim milk), while both the mother and infant were healthy and at different infections during the course of breastfeeding. A variety of infections were sampled, including common cold, mastitis and cracked nipples, ear infection, and gastrointestinal infection. Furthermore, leukocyte numbers as well as leukocyte subsets were determined using flow cytometry to explore associations between the leukocyte composition of HM and its miRNA content. In accordance with previous studies, [6,13] strong leukocyte responses to infection of the mother and/or the infant were recorded. Interestingly, specific leukocyte subsets were characteristic of different infection types. At the same time, miRNA responses to infection as well as associations with the leukocytic response were observed, supporting the notion that these HM molecules participate together with HM leukocytes in the immune protection of the mammary gland and the infant.

Associations with infection were first explored for total cell numbers, total RNA content, fat content, and leukocytic content and composition of HM. The total HM cell count (also referred to as milk somatic cell count in animal studies) [65] was not different between the healthy and the common cold cohorts (p=0.38), which is in agreement with previous literature [6]. However, there was a tendency of significant difference in the total cell number between the healthy cohort and the other infection cohort, which included mastitis and cracked nipples (amongst and other infections) (p=0.07), potentially due to mastitis, which is known to be associated with increased total cell counts in HM [6] as well as in the milk of other mammals [65]. Further, HM fat content was found to be lower in mother/infant dyads with common cold (p=0.045). Considering the use of HM fat content as an indicator of breast fullness [66–68], this finding suggests that participants with common cold had fuller breasts when they donated the milk sample. This may be potentially reflective of a temporary reduction in breastfeeding due to maternal sickness, resulting in fuller breasts. In contrast, RNA content obtained from HM cells or fat was not associated with infection (p>0.05). Previous studies have also reported no associations of HM RNA content with different maternal and/or infant factors, such as the stage of lactation or milk removal [28,29]. As expected, and in accordance with previous literature [6], any infection of the mother and/or infant was strongly associated with an increase in the relative percentage of leukocytes in HM (p<0.05). For example, the common cold resulted in an average of total leukocyte percentage of 22.62 ± 25.38 % compared to 1.68 ± 0.96 % recorded for HM from healthy dyads. The total HM cell count per mL of HM was related to the percentage of leukocytes (p=0.013), showing that the infections in mammary gland is increasing the total human milk cells by accelerating the production of total immune cells.

To explore any responses of HM miRNAs to infection, a subgroup of cell and fat samples was sequenced using Solexa, with 441,356,898 reads obtained out of 770,374,554 total reads mapped to miRBase 21.0. The separate analysis of the fat and cell fractions of HM enabled us to examine differences between HM fractions and responses to infection. HM cells had more miRNA species compared to HM fat, which is in agreement with previous studies in both human and animal milk[28,29,47,69–72]. In turn, milk fat is known to be richer in miRNA species that skim milk or exosomes [22,32,73]. Collectively, HM is shown to be one of the richest sources of miRNAs in humans.

A total of 1,780 mature known miRNAs were determined **(Supplementary Table S1)**, which is the greatest number of known miRNAs profiled so far in HM. Although in our previous HM miRNA profiling studies [28,29], the same version of miRBase (21.0) was used, here more known miRNA species were detected, potentially due to the influence of infection, which slightly increased the HM miRNA species number, though this was not statistically significant (p= >0.05). Interestingly, specific known miRNA species were detected in HM only during infection, suggesting a specific targeted function of these miRNAs during infection. Specifically, the top 20 miRNAs in both HM fractions (cells and fat) and cohorts (healthy and infection) contributed 88.3% of all the detected known miRNAs. Some of the top miRNAs expressed were different between the healthy and infection cohorts, indicating specific roles of these miRNA species during infection. Similar to our previous studies [28,29], all the members of the let-7 family were expressed at high levels and most of them were ranked in the top 50 miRNAs. Although the total number of known miRNA species was not statistically different between the healthy and infection cohorts (p= >0.05), samples with high number of B or T lymphocytes contained more known miRNA species (p=0.027 and p=0.039, respectively), suggesting that increased number of T and B cells associated with infection may stimulate expression of specific immune-related miRNAs.

Further, approximately a quarter of the identified known miRNAs were differentially expressed between the healthy and infection cohorts. Of these, 70 known miRNAs were strongly differentially expressed with infection **(Supplementary Table S3)**. Many of these are known to play roles in immunity in various mammals. For example, miR-223-3p participates in regulating differentiation and proliferation of neutrophils [19]. miR-150-5p is a significant regulator of the Toll-like receptor (TLR), which is involved in both innate and adaptive immune responses [74]. miR-150-5p targets CXCR4, which is a modulator of B cell development and induces humoral immunity [75]. In addition, miR-155-5p was highly upregulated in the infection cohort, with this miRNA known to play important functions in immune system regulation, especially in lymphocytes and dendritic cells [76]. Further, it has been previously reported to regulate the innate immune system during mastitis caused by *Streptococcus uberis* in the dairy cow [77]. Some of the differentially expressed miRNAs in HM were also homologous with bovine milk miRNAs in the presence of mammary gland inflammation, such as mastitis, including miR-150-5p [34,43], miR- 223-3p [33,34,43], miR-148 [43], miR-142-5p [33,34], miR-142-5p/3p, hsa-miR-132-3p, miR-455-5p, miR-139-5p, miR-503-5p, miR-424-5p, miR-450b, bta-miR-450a-5p, miR-31-5p, miR-145-5p [43], miR-338-3p, hsa-miR-339-5p (bovine mature name: bta-miR-339a), miR-34a-5p, and miR-338-3p [34]. The vast majority of the highly differentially expressed miRNAs (62 out of 70) were upregulated during infection compared to when both the mother and infant were healthy, supporting their specific involvement in the protection of the mammary gland and infant and facilitation of recovery.

In addition to known miRNA species, previous studies have recently revealed HM as a rich source of novel miRNA species [22,28,29]. Although 151,801,767 total reads were unannotated to all RNA categories, only 65,878 reads matched the strict novel miRNA criteria used here, and this translated to 592 novel mature miRNA species **(Supplementary Table S2)**. Of these, high-confidence novel miRNAs were determined as those identified in more than 2 samples with more than 20 reads [22,29]. Therefore, 95 novel miRNAs matched the high-confidence criterion. The number of discovered novel miRNA species was not related between cell and fat fractions, or between healthy and infection cohorts (both p>0.05). This was consistent with normal maternal factors, explaining that the novel miRNAs need to be first validated by other technologies to be reliably used for statistical analysis [22,28].

For further qRT-PCR validation of the sequencing results, seven miRNAs (let-7f- 5p; miR-1levels and148a-3p; miR-181a5p; miR-182-5p; miR-375; miR-99b-5p) thought to be involved in immunity and immune system regulation **(Supplementary Table S11)** were selected based on their expression patterns and differential expression levels, and were tested not only in HM cells and fat, but also in skim milk. These miRNAs are known to be highly expressed in HM [28–30] and they were found to be differentially expressed between the healthy and infection cohorts **(Supplementary Table S11)**. In a comparison of the different HM fractions, skim milk showed lower expression of miRNAs compared to cells and fat, whereas fat was lower compared to cells **(Figure 5)**. Recent studies comparing the three milk fractions in the dairy cow reported that milk cells had more miRNA species than fat, and the latter was higher than skim milk [22,28,78]. Interestingly, we followed a participant with mastitis during different stages of the infection (from the initiation of the infection to the recovery period during three weeks of infection). All the seven miRNAs tested here were expressed at low level in the HM cell fraction at the beginning of mastitis (stage 1) **(Figure 6A)**, but were then upregulated in both subsequent stages (full infection and during decline of infection stage), suggesting the potential use of these miRNAs as diagnostic markers and indicators of disease progression as well as response to treatment. In contrast, 5 of the tested miRNAs (let-7f-5p; miR-181a5p; miR-182-5p; miR-375; miR-99b-5p) showed high expression in mastitis stage 1 in HM fat, but were then downregulated in progressing mastitis **(Figure 6B)**. These differences between HM fractions likely reflect the lactocyte-specific character of the milk fat and demonstrate the need to carefully consider specific milk miRNAs during milk fractionation for use as biomarkers of disease, the health status of the lactating mammary gland, or lactation performance [78].

To determine the biological and cellular functions of the targets of the miRNA species discovered here, the highly expressed or differentially expressed miRNAs were used in a series of prediction tools. The majority of the targets of the top 50 more abundant miRNAs and the top 50 differentially expressed miRNAs were same. These targets (genes) were found to participate in immune system and immune response **(Supplementary Table S14-S15)**. Targets of the most abundant miRNAs in HM were associated with response pathways of the immune system to *Staphylococcus aureus* **(Supplementary Table S15)**, which is thought to be primarily responsible for mastitis in women. It has been previous demonstrated that most of the top 20 miRNAs in HM are involved in development and growth [27], and also in the biosynthesis of different HM components, such as protein and lactose [29]. Here, highly expressed miRNAs were also found to have similar roles in development, especially in cellular differentiation and apoptosis, reinforcing the potential involvement of these miRNAs in milk synthesis and in the infant.

This study examined the effects of different types of infection of the mother and/or the infant during breastfeeding on the miRNA composition of HM and associations with HM leukocyte subsets. HM is a biofluid highly rich in miRNAs, especially in its cell and fat fractions, which rapidly respond to infection primarily by upregulation of their expression levels, similar to what has been previously shown and confirmed here for HM leukocytes [6,10,13]. With many known functions of these molecules in immune system and immunity, differentially expressed miRNAs in HM in response to infection may be used as biomarkers of the health status of the lactating mammary gland, particularly to follow disease progression and the response to treatment, as well as of lactation performance [15,22,79].

## ACKNOWLEDGMENTS

This publication was supported by an unrestricted research grant from Medela AG (Switzerland). The company had no input in designing or conducting the study or the decision to publish the manuscript. The gratitude is extended to The Geddes Hartmann Human Lactation Group at The University of Western Australia for the assistance. Many thanks are extended to all mothers who participated in this study, to the Australian Breastfeeding Association for assistance in recruitment of participants.

## AUTHOR CONTRIBUTIONS

All authors have contributed equally.

## SUPPLEMENTARY INFORMATION

Supplementary Figures S1-S4

Supplementary Tables S1-S15

